# Identification of conserved cross-species B-cell linear epitopes in human malaria: A subtractive proteomics and immuno-informatics approach targeting merozoite stage proteins

**DOI:** 10.1101/2023.11.23.568458

**Authors:** Sebastian D. Musundi, Jesse Gitaka, Bernard N. Kanoi

**Author notes:** Correspondence: Centre for Malaria Elimination, Institute of Tropical Medicine, Mount Kenya University, Thika, Kenya. Bernard N. Kanoi.

## Abstract

Human malaria, caused by five Plasmodium species (*P. falciparum, P. vivax, P. malariae, P. ovale*, and *P. knowlesi*), remains a significant global health burden. While most interventions target *P. falciparum*, the species associated with high mortality rates and severe clinical symptoms, non-falciparum species exhibit different transmission dynamics, remain hugely neglected, and pose a significant challenge to malaria elimination efforts. Recent studies have reported the presence of antigens associated with cross-protective immunity, which can potentially disrupt the transmission of various Plasmodium species. With the sequencing of the Plasmodium genome and the development of immunoinformatic tools, in this study, we sought to exploit the evolutionary history of Plasmodium species to identify conserved cross-species B-cell linear epitopes in merozoite proteins. We retrieved Plasmodium proteomes associated with human malaria and applied a subtractive proteomics approach focusing on merozoite stage proteins. Bepipred 2.0 and Epidope were used to predict B-cell linear epitopes using *P. falciparum* as the reference species. The predictions were further compared against human and non-falciparum databases and their antigenicity, toxicity, and allergenicity assessed. Subsequently, epitope conservation was carried out using locally sequenced isolates from a malaria-endemic region in western Kenya (n=27) and Kenyan isolates from MalariaGEN version 6 (n=131). Finally, physiochemical characteristics and tertiary structure of the B-cell linear epitopes were determined. The analysis revealed eight epitopes that showed high similarity (70-100%) between falciparum and non-falciparum species. These epitopes were highly conserved when assessed across local isolates and those from the MalariaGEN database and showed desirable physiochemical properties. Our results show the presence of conserved cross-species B-cell linear epitopes that could aid in targeting multiple Plasmodium species. Nevertheless, validating their efficacy *in-vitro* and *in-vivo* experimentally is essential.

## Introduction

Malaria remains a significant health burden in Sub-Saharan Africa, especially for children under five years and pregnant women. Among the five Plasmodium species (*P. falciparum, P. vivax, P. ovale, P. malariae*, and the zoonotic *P. knowlesi)* that cause human malaria, *P. falciparum* accounts for the highest number of malaria cases globally while *P. vivax* shows a broader geographical distribution (Price et al., 2020; World Health Organization, 2022). While significant progress has been made in malaria control efforts over the past two decades, the focus is primarily on *P. falciparum* due to its clinical severity. Less emphasis is placed on non-falciparum species despite evidence of existing differing transmission patterns with *P. falciparum*. For instance, despite a decline in *P. falciparum* malaria cases in Tanzania, persistent transmission of *P. ovale* and *P. malariae* between 2010 and 2016 continued to account for the increased number of malaria cases (Yman et al., 2019). Similarly, *P. knowlesi* emerged as the primary cause of malaria in Sabah Malaysia after a significant decline in cases due to *P. vivax* and *P. falciparum* infections (William et al., 2013; Cooper et al., 2020). The differences in transmission in non-falciparum species may explained by the presence of hypnozoite stages (Collins and Jeffery, 2005), differing seasonality and asymptomatic infections (Tarimo et al., 2022), and differences in gametocytogenesis (Holzschuh et al., 2022). To successfully eliminate malaria, it is crucial to implement strategies targeting all Plasmodium species linked with human malaria to improve the health outcomes of vulnerable populations globally.

The evolutionary relationship between Plasmodium species is a valuable resource for developing interventions targeting multiple species. This approach has already been exploited to identify potential vaccine targets for *P. vivax* from *P. falciparum* homologs (Duffy and Patrick Gorres, 2020). Since Plasmodium species emerged and evolved from a common ancestor, the structure and function of some proteins remained similar, increasing the possibility of homologous proteins recognizing similar B-cell linear epitopes during infections. Existing evidence already points out cross-reactivity between homologous proteins in Plasmodium. For example, recombinant *P. vivax* apical asparagine (Asn)-rich (PvAARP) was strongly recognized by *P. knowlesi* patient sera, cross-reacted with the apical side of the *P. knowlesi* merozoites and inhibited erythrocyte invasion by *P. knowlesi* with the mechanism linked to similar B-cell epitopes in both species (Muh et al., 2018). Additionally, functionally, non-reciprocal cross-species immunity between *P. vivax* Duffy Binding Protein (PvDBP), a protein mediating reticulocyte invasion, and *P. falciparum* VAR2CSA, a protein involved in parasite sequestration in the placenta has been observed (Gnidehou et al., 2019; Mitran et al., 2019). VAR2CSA and PvDBP are not functionally related but share a Duffy binding-like domain, with cross-reactive epitopes in *P. vivax* blocking the adhesion of VAR2CSA to its placental receptor, chondroitin Sulfate A (Gnidehou et al., 2019; Mitran et al., 2019). Antibodies arising from PvDBP cross react with the DBL3X region of VAR2CSA through a 34 amino acid conserved region and have also been shown to elicit adhesion binding antibodies against CSA, suggesting their role as a potential vaccine candidate (Iyamu et al., 2023). Thus, by leveraging the evolutionary relationship and shared epitopes among Plasmodium species, interventions that provide broad protection against different malaria-causing parasites can be developed, thereby contributing to global efforts aimed at malaria elimination.

The sequencing of the Plasmodium genomes in the post-genomic era has greatly accelerated the discovery of potential vaccine candidates. When coupled with subtractive proteomics, which seeks to subtract pathogen targets needed for the organism’s survival but not in the host, immuno-informatics, which predicts the presence of epitopes, reduces the time required to identify antigen targets for experimental validation. Herein, we utilized a subtractive proteomics and immunoinformatic approach to identify homologous proteins present in merozoites in *P. falciparum*. Cross-species B-cell linear epitopes sharing high similarity with *P. falciparum* were identified, and their conservation was assessed using locally sequenced isolates from a malaria endemic region and those deposited in MalariaGEN version 6. We identified eight conserved cross-species B-cell linear epitopes that need further assessment as potential vaccine targets that can be developed for falciparum and non-falciparum malaria as part of the consented efforts to control and eliminate the scourge.

### Materials and Methods Removal of redundant proteins

The workflow applied in identification of conserved B-cell linear epitopes is summarized in Figure 1. Initially, the latest complete proteome sequences for *P. falciparum* 3D7, *P. vivax* P01, *P. malariae* UG01, *P. ovale* GH01, *P. knowlesi* Strain H were downloaded from PlasmoDB version 63, and the human proteome from NCBI. CD-HIT was used to remove redundant protein sequences using a threshold of 90% (Fu et al., 2012). After that, the *P. falciparum* proteome was used as the reference for downstream analysis.

**Figure 1.**
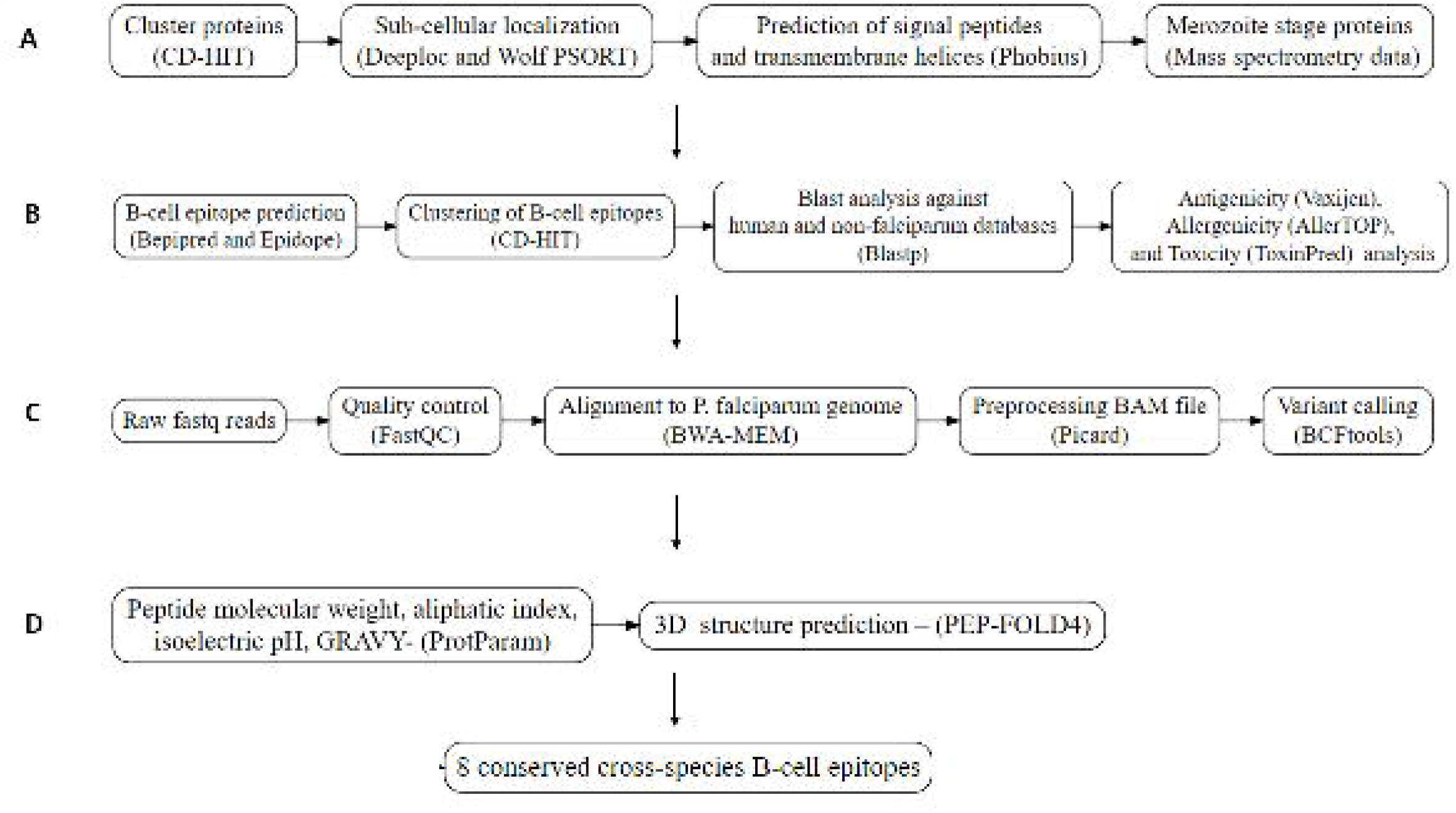
Workflow utilized for the prediction of conserved B-cell linear epitopes. (A). Subtractive proteomics. (B) Epitope prediction and analysis. C). Conservation analysis from merozoite protein from local Kenyan isolates (n=27) and MalariaGEN (n=131). D). Structural and physicochemical properties.

### Subcellular localization and removal of intracellular proteins

WoLF PSORT and Deeploc were employed to predict the subcellular location of the selected proteins. In WoLF PSORT, amino acid sequences are converted to numerical localization features based on their sorting signals, functional motifs, and amino acid composition. Then, the nearest neighbor classifier is used to predict the location of proteins in 10 different sites (Horton et al., 2007). Deeploc predicts utilizing a deep learning approach by using datasets from eukaryotic and human proteins (Thumuluri et al., 2022). Intracellular proteins, such as those in the cytoplasm and nucleus, were excluded from further analysis. However, proteins present in *P. falciparum*-specific membrane-bound organelles such as rhoptries or micronemes cannot be accurately classified by WoLF PSORT and Deeploc since these organelles are absent from their predictions. To cater to proteins in the rhoptries and micronemes, membrane-bound organelles with similar properties to these plasmodium organelles were selected. Afterward, proteins found in intracellular organelles such as the mitochondria, ribosomes, and apicoplast were removed using specific keywords in their protein names. Transcription, translation, RNA, and DNA binding proteins were eliminated using the same approach. Lastly, enzymes were also excluded from the analysis.

### Prediction of surface proteins

The presence of a signal peptide and transmembrane helix was assessed to narrow down the proteins of interest. Signal peptides (SPs) are short peptide sequences majorly present on the N-terminus of protein sequences destined towards secretory cells or on the cell membrane. At the same time, transmembrane helices are membrane-bound proteins that play various roles, including anchoring transmembrane proteins to the cell membrane (Krogh et al., 2001; Owji et al., 2018). Phobius was used to predict the presence of a signal peptide and transmembrane helix. Phobius utilizes hidden Markov models with sub-models capable of differentiating between signal peptides and transmembrane helices (Käll et al., 2004). Proteins with either one signal peptide or transmembrane helix were selected for further analysis.

### Merozoite protein selection

Malaria infections begin when sporozoites are injected into the bloodstream through a bite by an infected female Anopheles mosquito. The sporozoites migrate to the liver where they infect hepatocytes, and within 7-10 days, the parasite develops to become merozoites released into the bloodstream to infect red blood cells, initiating the intra-erythrocytic stage of the disease. The replication cycle in the blood stage takes approximately 24 hours in *P. knowlesi*, 48 hours in *P. falciparum, P. ovale, and P. vivax, and* 72 hours in *P. malariae*. This asexual stage is associated with clinical manifestations of malaria. Intra-erythrocytic stage antigens are presented on the surface and in the apical organelles of merozoites, and on the surface of infected red blood cells where come into direct contact with the host immunity this makes these proteins targets for naturally acquired immunity as well as anti-disease vaccine targets. Thus, mining of mass spectrometry data from PlasmoDB was performed to identify proteins expressed in the merozoite stage (Supplementary Table 1).

### B-cell linear epitope prediction

B-cell linear epitopes are specific regions within the primary structure of proteins that interact with antibodies. The presence of B-cell linear epitopes was predicted using Bepipred 2.0 and Epidope using their default settings. Bepipred 2.0 utilizes a random forest algorithm which has been trained on epitopes annotated from 649 antigen-antibody structures derived from the Protein Data Bank (PDB), while Epidope utilizes deep neural networks using data derived from Immune Epitope Database (IEDB) (Jespersen et al., 2017; Collatz et al., 2021). The resulting predictions were then converted into FASTA format, where each entry in the header consisted of the indexed Plasmodium protein from the database, along with the corresponding start and end positions of the predicted epitopes separated by a colon and underscore for Bepipred and Epidope, respectively. The sequence of the predicted epitopes was included in the second line of each entry to complete the FASTA file. The predicted B-cell linear epitopes were clustered together using CD-HIT to identify common epitopes using a similarity of threshold 0.9. The start and end positions of the predicted B-cell linear epitopes were individually visualized using Protter to ensure they were not present in the signal peptide or transmembrane domain region (Omasits et al., 2014).

### Homology analysis

The predicted B-cell linear epitopes from *P. falciparum* were blasted against an indexed non-falciparum database comprising of the CD-HIT clustered proteins from *P. ovale, P. malariae, P. knowlesi*, and *P. vivax* and human proteome using an e-value of 1e-05. *P. falciparum* B-cell linear epitopes that showed similarities with human proteins can induce autoimmune reactions or downregulate the human response and were removed from further analysis. The remaining cross-species epitopes of length between 10-30 amino acids with a percentage identity >=70 and query coverage >=95 % were selected for further analysis.

### Antigenicity, Toxicity, and Allergenicity

The antigenicity, toxicity, and allergenicity of *P. falciparum* B-cell linear epitope sequences were evaluated. Vaxijen, a tool that allows for antigenicity prediction based on the physicochemical properties of proteins without relying on sequence alignment, was employed to determine the antigenicity of the selected peptides. Peptides were considered antigenic if the obtained score was greater than 0.5 for parasites (Doytchinova and Flower, 2007). Toxinpred2 was used to assess peptide toxicity, while AllergenPro was employed to evaluate allergenicity. ToxinPred2 classifies toxins based on their BLAST similarity with existing known toxic peptides, similar motifs, and prediction models (Sharma et al., 2022). AllergenPro compares peptide sequences with known allergen sequences and motifs from NCBI PubMed, UniProtKB/ Swiss-Prot, and IUIS allergen databases (Kim et al., 2009). B-cell linear epitopes that were antigenic, non-toxic, and non-allergens were selected for the next step.

### Conservation Analysis

Parasite DNA extracted from whole blood samples (n=27) cultured in vitro until the schizont-merozoite stages from a surveillance study in four selected islands in Lake Victoria between July 2014 and July 2016, together with MalariaGen data version 6 from Kenyan isolates was used to assess the conservation of the predicted B-cell linear epitopes. The previously published study was approved by the Mount Kenya University Ethics Review Committee (038/2014) and Kenyatta National Hospital - University of Nairobi (KNH-UoN) ethical review committee (P609/10/2014), and the sequencing data was used here as secondary analysis (Gitaka et al., 2017). For bioinformatics analysis, the quality of paired-end reads was assessed using FastQC (Leggett et al., 2013). The *Plasmodium falciparum* genome was retrieved from PlasmoDB and was indexed using the Burrows-Wheeler Algorithm (BWA) (Li and Durbin, 2010). The paired-end reads were mapped to the index genome to generate a SAM file. Samtools was used to convert the generated SAM file to BAM format. After sorting and indexing the BAM file, Picard was used to preprocess the files before variant calling. Bcftools utilities were used to call, normalize, and filter variants in the protein containing the B-cell linear epitope based on a haploid model, including those with a base quality score > 20 and minimum read depth >=5. Bcftools consensus was then used to replace specific positions in the reference where variants occurred for each sample (Danecek et al., 2021). The generated FASTA files were translated with their 3D7 reference PlasmoDB and aligned in MUSCLE (Edgar, 2004). The aligned protein file was downloaded, and the regions corresponding to the predicted epitope of interest were retrieved using an in-house bash script. The resulting conservation logos were plotted using WebLogo.

### Structural and Physiochemical properties

The PEP-Fold4 server was used to predict the tertiary structure of peptides with lengths of 5-50 amino acids (Rey et al., 2023). ProtoParam from Expasy assessed the physiochemical properties of conserved B-cell linear epitopes. The parameters evaluated by ProtoParam included molecular weight, instability index, aliphatic index, isoelectric pH (pI), and grand average of hydropathicity index (GRAVY)(Wilkins et al., 1999). The instability index estimates the protein stability in a test tube. An instability index greater than 40 is regarded as unstable, while a value below 40 indicates the peptide is stable. The aliphatic index refers to the relative volume occupied by aliphatic side chains and is regarded as a key factor in increasing the thermostability of globular proteins. High aliphatic index proteins tend to be more thermally stable. GRAVY indicates the sum of hydropathy values of all amino acids divided by all residues present. Negative GRAVY scores indicate the peptide is polar, while positive GRAVY scores indicate the protein is non-polar (Doytchinova and Flower, 2007).

## Results

### Removal of redundant proteins

The whole proteomes of the five Plasmodium species were passed through a subtractive proteomics pipeline. CD-HIT refined each proteome, removing redundant proteins at the 90% threshold. The *P. ovale* proteome had the highest number of proteins after removing paralogs, while *P. malariae* had the greatest number of paralogs (Supplementary Table 2).

### Subcellular classification

The cellular location of the remaining 5289 proteins from *P. falciparum* was carried out using Deeploc and WoLF PSORT. The total number of proteins present in the intracellular, extracellular, and membrane-bound regions for Deeploc was 2637, 1005, and 1628, while those for WoLF PSORT were 3188, 1299, and 807, respectively. A consensus approach was used to select proteins in the extracellular region and membrane-bound organelles from Deeploc and WoLF PSORT, resulting in 3488 proteins. Membrane-bound proteins were excluded using key search names, including those in the apicoplast, mitochondria, golgi body and endoplasmic reticulum. Similarly, enzymes, ribosomal proteins, eukaryotic transcription, and translation factors were excluded. In total, 1046 proteins were removed from the remaining proteins, leaving behind 2442 proteins.

### Surfaceome analysis and protein selection

Signal peptides and transmembrane helices are key features of proteins located on the extracellular surface. From the remaining 2442 proteins, Phobius predicted 412 proteins to either contain a signal peptide and one transmembrane helix or a signal peptide without a transmembrane helix. Since Phobius differentiates signal peptide and transmembrane helices, it predicted 711 proteins to have one transmembrane helix. A consensus approach was also applied, resulting in 953 proteins either containing a signal peptide or having one transmembrane helix. Down-selection of the 953 proteins against a list of published proteins in the merozoite stage reduced the total number of proteins to 366.

### B-cell epitope Analysis

Linear B-cell epitope prediction was done using the IEDB tool Bepipred 2.0 and Epidope on the 366 proteins. Bepipred 2.0 and Epidope generated 3827 and 1745 B-cell linear epitopes with a minimum length of 9 amino acids. Cluster analysis for the predicted epitopes using a threshold of 0.9 from CD-HIT generated 3637 clusters. Blast analysis for the predicted B-cell linear epitopes against non-falciparum species and human proteome revealed a total of 919 and 32 epitopes, respectively. The homologous human epitopes were excluded from further analysis due to their ability to induce auto-antibodies. Subsequent filtering of homologous non-falciparum regions, which showed a similarity >=95%, minimum coverage of 70%, and alignment length between 9-30 amino acids, resulted in 159 cross-species epitopes. Antigenicity analysis using Vaxijen 2.0 revealed the presence of 69 antigenic epitopes. Of these, 31 epitopes were predicted to be allergens from AllerTop, while 14 epitopes were predicted to be toxins from Toxinpred2.0 (Supplementary Table 3). Six of the remaining 24 epitopes were located either within the signal peptide region or within the intracellular region of transmembrane helix using Protter, resulting in a final 18 epitopes (Supplementary Table 4). B-cell linear epitopes in present in more than 3 *Plasmodium* species were further selected for conservation analysis. Additional information regarding *P. falciparum* antigen selected, start and stop position of the epitope, antigenicity, phenotype information from PhenoPlasm, and their presence in non-falciparum species was extracted (Table 1).

**Table 1.**
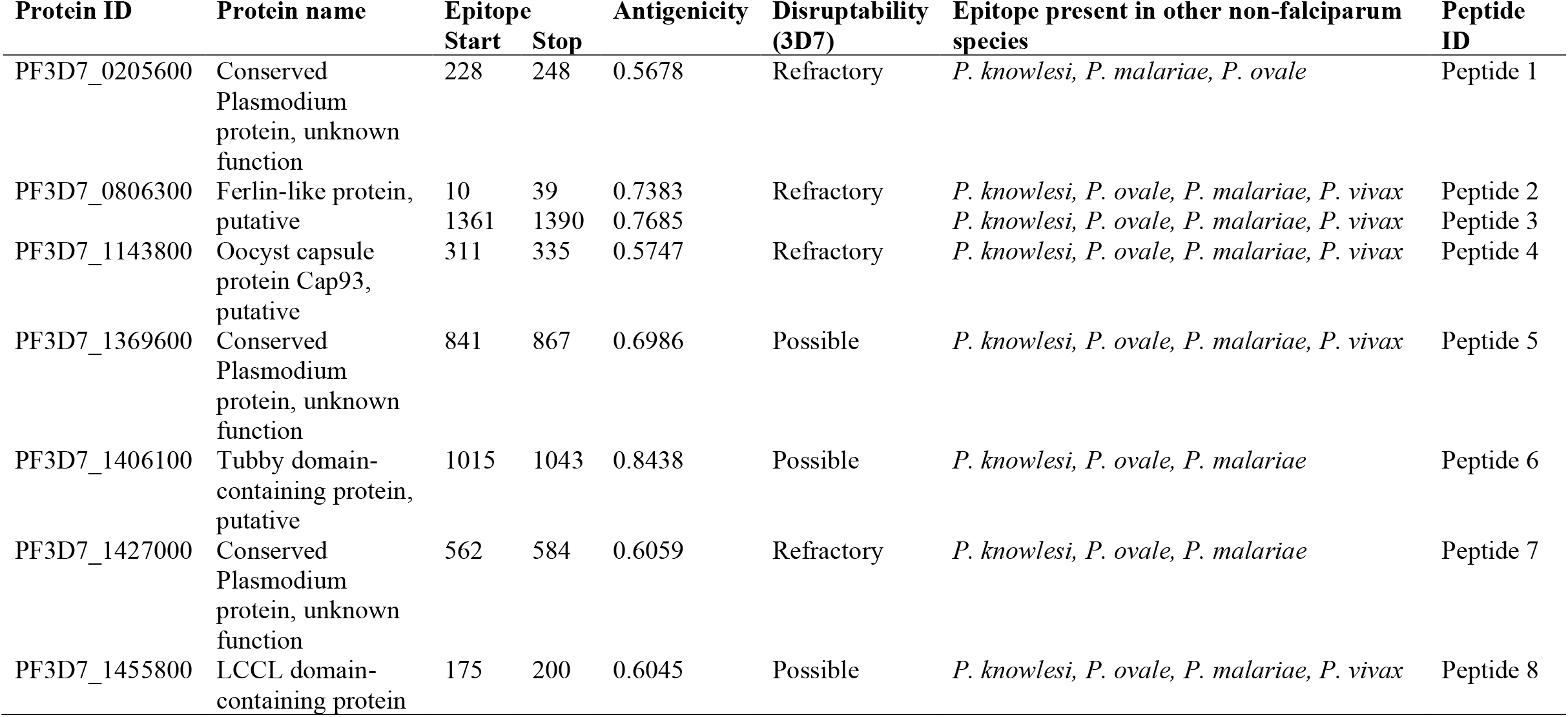
*Plasmodium falciparum* protein containing the selected B-cell linear epitopes, antigenicity scores, essentiality of the gene and presence in other non-falciparum species Complete list is shown is Supplementary Table 3.

Cross-species blast analysis of the predicted B-cell linear epitopes revealed the presence of 8 epitopes in more than four Plasmodium species based on the applied blast parameters (Supplementary Table 4). Some predicted B-cell linear epitopes showed high similarity between *P. falciparum* and one or two other Plasmodium species (Supplementary Table 5). The antigenicity score of the predicted peptides ranged from 0.5040 to 1.2264 (Table 1). Proteins, PF3D7_0205600, PF3D7_0806300, PF3D7_1143800, and PF3D7_1427000 containing the B-cell linear epitopes showed an essential role in the parasite) and were designated as refractory in Phenoplasm while the proteins PF3D7_1369600, PF3D7_1406100 and PF3D7_1455800 containing did not hinder parasite growth (Table 1). Peptides 2 and 4 showed the least changes in their amino acid sequences when plasmodium species were compared against the other non-falciparum species (Figure 2). The conservancy assessment of the eight peptides based on locally sequenced isolates in *P. falciparum* (n=27) and Kenyan isolates from MalariaGEN (n=131) showed high conservation with no change in the amino acid sequence being reported in the predicted region (Figure 2).

**Figure 2.**
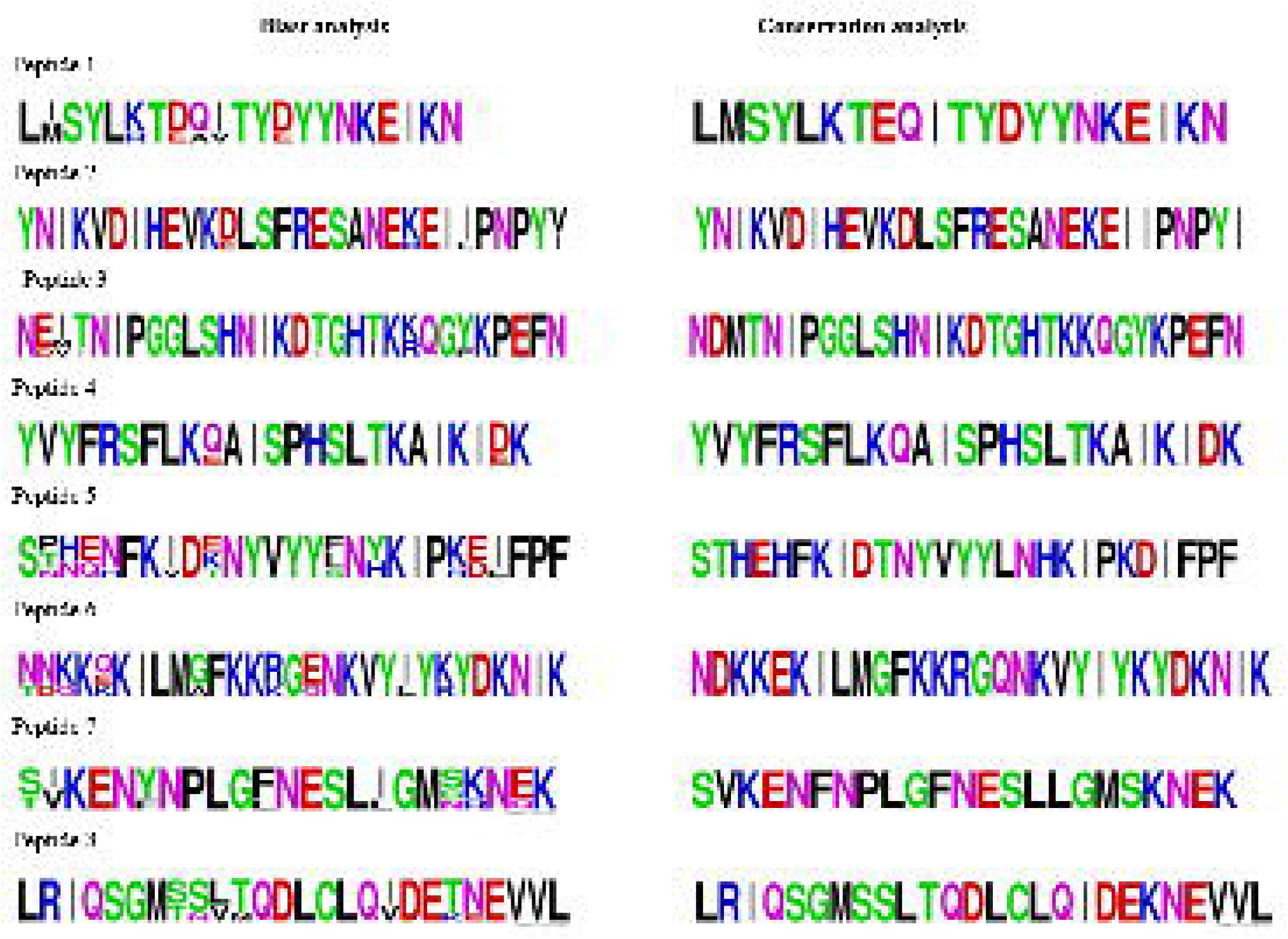
Blast and conservation analysis of the 8 predicted B-cell linear epitopes. Blast analysis incorporates predicted B-cell linear epitopes in falciparum against non-falciparum species with percentage similarity >70%, alignment length <=30 and coverage >=95. B-cell linear epitopes conservation analysis was against local sequenced isolates from Homabay (n=27) and MalariaGEN (n=131) from Kenya. The height of the amino acid sequence at each position represents its relative frequency in both the blast and conservation analysis.

### Structural and Physicochemical properties

The physicochemical properties of the final eight antigenic, non-toxic, and non-allergens epitopes were assessed. The properties, including the molecular weight, aliphatic index, pI, GRAVY, instability index, and total number of positively and negatively charged amino acids, are summarized in Table 2. The molecular weight of the eight selected peptides ranged from 2582.91 to 4813.77. The results also showed that all peptides were hydrophilic except peptide 8, while only peptide 1 was predicted to be unstable. The pI of the epitopes ranged from 4.32 to 10.22, while the most aliphatic and thermostable was peptide 8. The 3D prediction results from PEP-FOLD-4 revealed that peptides 1, 3, 5, and 7 had helices and coils, while peptides 2, 4, 6, and 8 comprised helices, strands, and coils. (Supplementary Figure 1).

**Table 2.**
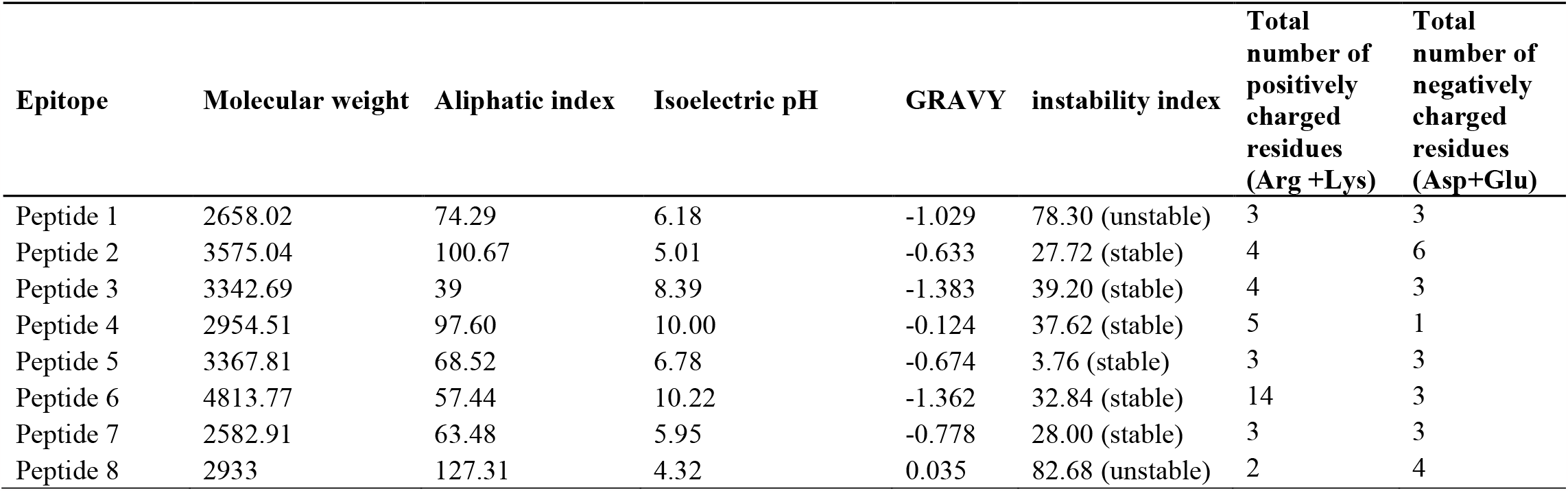
Predicted physicochemical properties of conserved B-cell linear epitopes.

## Discussion

Malaria, the most common tropical disease, remains a major threat to Sub-Saharan Africa. The different transmission patterns associated with Plasmodium species threaten malaria elimination efforts. Recent studies have documented that cross-protective immunity may play a key role in developing vaccines that may halt malaria transmission (Muh et al., 2018, 2020). Combining subtractive proteomics and immune-informatic approaches offers a faster identification of potential targets before experimental validation. The increase in high-throughput genomic sequencing of various malaria isolates across the globe also allows us to examine if the predicted targets are conserved, which is essential in vaccine development.

Prioritization of potential targets from the proteome level relies on subtractive proteomics to narrow down to a few essential proteins. The decision tree implemented followed commonly established procedures, including proteome clustering, subcellular localization, surfaceome predictions, and similarity analysis using BLAST (Aguttu et al., 2021). The use of CD-HIT ensured that only non-redundant proteins were assessed. Two tools, WoLF PSORT and Deeploc, were utilized to predict the subcellular location of the potential targets. Unlike bacterial organisms whose tools for determining subcellular locations are well established, a significant gap still exists in determining the subcellular location of protozoan parasites since they contain special organelles such as dense granules, rhoptries and micronemes. Thus, utilizing a consensus approach from two tools that predict the location using different approaches increases the chance of picking up more targets. A similar analysis carried out by Componovo et al. also revealed that following strict classification by only choosing extracellular and plasma membrane proteins also left out numerous proteins misclassified in the other organelles, which showed high immunoreactivity against the patient’s sera (Camponovo et al., 2020). Therefore, adding an extra step involving eliminating specific protein names found in intracellular organelles not involved in the invasion process improves the subcellular classification of proteins.

Blood-stage antigens such as those on merozoites are essential for red blood cells invasion, hence excellent vaccine candidates. Ideal vaccine candidates, especially in malaria, should possess low levels of polymorphism and be highly antigenic (Ouattara et al., 2015). Our results identified eight B-cell linear epitopes, which were predicted to be antigenic and showed high levels of conservation, especially across *P. falciparum* isolates. In addition, the isolates showed high similarity with other non-falciparum species linear epitope regions. The conserved nature of the epitopes and their high similarity with non-falciparum species indicates the likelihood of cross-species reactivity and immunogenicity, although this needs to be experimentally validated. *P. falciparum* antigen VAR2CSA and *P. vivax* DBP (Gnidehou et al., 2019), Pfs48/45 and Pvs48/45 (Bansal et al., 2017), and *P. vivax* AMA1, and *P. falciparum* AMA1 (Igonet et al., 2007) have shown cross-species reactivity. Cross-protective immunity towards human malaria has also been shown experimentally between *P. vivax* AARP and *P. knowlesi* AARP and *P. vivax* RhopH2 and *P. knowlesi* RhopH2 (Muh et al., 2018; Ahmed et al., 2022). Furthermore, the presence of conserved B-cell epitopes between *P. vivax* RhopH2 and *P. knowlesi* RhopH2 has already been established (Ahmed et al., 2022). Experiments in mice models utilizing Pvs48/45 and Pfs48/45 antigens also showed the ability of the two antigens to cross-boost each other (Cao et al., 2016). In natural populations, mixed infections of various species occur, increasing the possibility of inducing cross-protection against multiple species. In the current context, we postulate an immune response targeting multiple different parasite species, driven by their shared B-cell epitopes. Additionally, we anticipate a reinforced secondary immune response upon repeated exposure to either parasites species.

A stable structure among other physiochemical properties are key for a novel vaccine candidate. Here, seven epitopes were identified and predicted to have a stable structure, while one epitope had an unstable structure. Cyclization of the peptide sequence is one of the approaches that could be applied to make the peptides more stable before further downstream analysis (Hamley, 2022). Furthermore, epitopes were shown to have an isoelectric pH (pI) ranging from 4.32 to 10.22. The index pI is an important factor in selecting peptide vaccine since, at their pI, they have no electric charge, and considering that B-cell receptors prefer charged and polar antigen residues, their immunogenicity tend to decrease (Wang et al., 2018). Therefore, it ideal for the pI to not fall within the range of human body tissues (pH 7.2 to 7.6) as in the case of all epitopes. All these parameters suggest these epitopes are appropriate candidates for in-vitro and vivo validation experiments.

In conclusion, in this study, we utilized subtractive proteomics and immunoinformatics techniques to identify conserved cross-species B-cell linear epitopes in the merozoite stage. These epitopes are antigenic, non-toxic, non-similar to humans, non-allergens, and have suitable physiochemical properties. Evaluation of the identified conserved B-cell epitopes could contribute to development of vaccines that induce cross-protective immunity. Nevertheless, there is a need to confirm the functionality of the individual conserved epitopes in vivo and in-vitro experiments.

## Author Contributions

BK and JG conceived the idea. JG cultured malaria parasites and coordinated samples for sequencing. SM developed the methodology and analyzed the bioinformatic data. SM wrote the first draft of the manuscript. BK and JG revised the manuscript. The final manuscript was read and approved by all authors.

## Acknowledgment

We wish to acknowledge the Mount Kenya University research team for their insights into this work.

## Data Availability Statement

The datasets presented in this study has been deposited in DDBJ under the BioProject Accession Number PRJDB12148 and other publicly available datasets from MalariaGEN.

