## Supplementary figures and images for "Identification of conserved cross-species B-cell linear epitopes in human malaria: A subtractive proteomics and immuno-informatics approach targeting merozoite stage proteins"

### Supplementary Figure 1.PNG

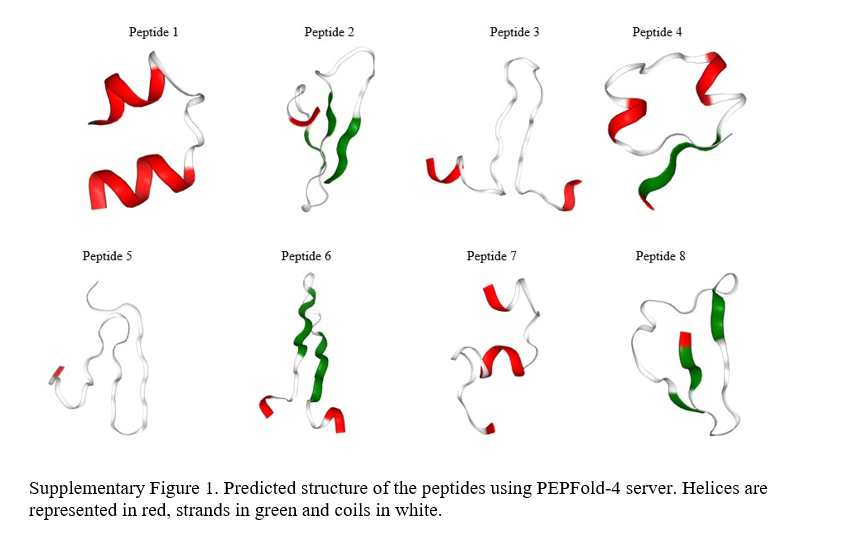
